# Exact p-values for large-scale single step genome-wide association, with an application for birth weight in American Angus

**DOI:** 10.1101/555243

**Authors:** Ignacio Aguilar, Andres Legarra, Fernando Cardoso, Yutaka Masuda, Daniela Lourenco, Ignacy Misztal

## Abstract

**BACKGROUND:** Single Step GBLUP (SSGBLUP) is the most comprehensive method for genomic prediction. Point estimates of marker effects from SSGBLUP are often used for Genome Wide Association Studies (GWAS) without a formal framework of hypothesis testing. Our objective was to implement p-values for GWAS studies in the ssGBLUP framework, showing algorithms, computational procedures, and an application to a large beef cattle population.

**METHODS:** P-values were obtained based on the prediction error (co)variance for SNP, which uses the inverse of the coefficient matrix and formulas to compute SNP effects.

**RESULTS:** Computation of p-values took a negligible time for a dataset with almost 2 million animals in the pedigree and 1424 genotyped sires, and no inflation was observed. The SNP passing the Bonferroni threshold of 5.9 in the −*log*10 scale were the same as those that explained the highest proportion of additive genetic variance, but the latter was penalized (as GWAS signal) by low allele frequency.

**CONCLUSION:** The exact p-value for SSGWAS is a very general and efficient strategy for QTL detection and testing. It can be used in complex data sets such as used in animal breeding, where only a proportion of pedigreed animals are genotyped.

## INTRODUCTION

Detection and mapping of causal genes and QTLs in livestock genetics is usually done by Genome-Wide Association Studies (GWAS). The most frequent GWAS method is the single-marker fixed regression, where one marker is fit at a time in the model, and correction for polygenic genetic effects is made using a relationship matrix. This is also known as EMMAX [1–3]. However, use of EMMAX implies that genotyped animals have phenotype and vice versa. In livestock, this requirement is impossible for animals such as dairy bulls that do not have female-related phenotypes (e.g. milk yield). In general, many animals that have own phenotype (e.g. growth) would benefit from phenotypic information from relatives (e.g. growth in daughters, ancestors and collaterals). Typically, phenotypes of related, animals are “projected” into genotyped animals [4–6] in a process called de-regression, which has been successfully used to detect and map QTLs [7,8].

De-regression is a cumbersome process, usually not optimal because it is an approximation that lost information and can lead to inaccuracies due to unaccounted for selection, or common effects that are just ignored. In particular using de-regression leads to double counting when the genotyped population includes both sires and their progeny. Legarra and Vitezica [9] showed that an extension of the single-marker EMMAX regression is a two-trait variance component model, where the two traits are actual phenotype and “gene content” at the marker. However, the method would be very slow for a whole genome scan because it requires maximization of a REML likelihood at each marker.

The GBLUP or SNP-BLUP framework [10–12] allows for the joint estimation of marker effects and the automatic correction of genetic structure in the population. The so-called single step methods (i.e., Single Step GBLUP (SSGBLUP) and Single Step SNP-BLUP (SSSNP-BLUP)) project enotypes into phenotyped individuals, using pedigree relationships [13–16]. These “single step methods” allow the estimation of breeding values and also marker effects [15,17,18], and the latter have been used for GWAS analysis [19–21], typically by size of estimated marker effects or a similar measure such as the proportion of genetic variance explained by a marker or segment. A common pitfall of GWAS methods based on size of effects or variances explained is the lack of a formal framework for acceptance/rejection of hypothesis – in particular, there is no clearly defined statistic to be used for hypothesis testing, and there is no empirical statistical law or closed-form solution under the null hypothesis. The most common form of declaring significance is to use a threshold on explained genetic variance (but: how much?) by a set of contiguous markers (but: how many?) [22]. For instance, [19] studied “the 20 largest explanatory loci”, [20] studied “the 10 windows explaining the largest amount of genomic variance for gene annotation, gene network and pathway analyses” whereas [21] considered “1.5 Mb SNP windows that explained more than 0.50 % of the genetic variance”. Thus, there is no formal hypothesis testing. Although using estimates of markers effects considers correctly the *magnitude* of the estimated marker effects, it does not always consider correctly the *uncertainty* in the estimation of marker effects. Even worse, the use of different iterative schemes or the definition of “windows” of adjacent SNPs can be very arbitrary. This renders interpretation of signals more difficult‥

Recently, equivalences between GBLUP and single marker GWAS results have been shown in a series of papers [23–25] Namely, the statistic of single marker GWAS is equivalent to the statistic obtained from the marker estimate from GBLUP. In both cases the statistic used is 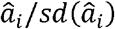 (i.e., marker effect estimate over its standard deviation). Although in EMMAX the marker is a fixed effect and in GBLUP it is random, a series of papers [23–25] at the statistic is the same in both methods. An application and comparison with Bayesian methods was shown in a data set of gait in horses [26]. Still, the method could not be applied to data sets consisting of mixtures of genotyped and ungenotyped animals.

Fortunately, Lu et al. [27] showed that the same logic used in [23–25] can be used in a SSGBLUP context. Simply stated, to obtain a statistical test for the effect of a single marker, we only need estimates of the breeding values of genotyped animals and their sampling distribution, and this can be readily obtained from SSGBLUP. Unfortunately, the article of Lu et al. [27] seems largely unnoticed because it focuses on feed efficiency and uses a small data set (i.e., 7,000 phenotyped animals from which 5,000 are genotyped), which does not show computational tools, challenges and bottlenecks.

In the present work, we show the implementation of Single Step GWAS (SSGWAS) with exact p-values as in [27], together with an application in a very large beef cattle data set. We describe algorithms and computational procedures along with their bottlenecks. As result of the data analysis, we find highly significant signals in loci already described by the literature, and good empirical behavior of the method.

## Methods

### Theory

The classical GWAS method (EMMAX) for marker *i* uses a linear model ***y*** = ***Xb*** + ***z***_*i*_*a*_*i*_ + ***u*** + ***e*** where vector ***b*** contains fixed effects, ***z***_*i*_ is a vector with “gene contents” (0, 1, or 2 at the marker), *a*_*i*_ is the substitution effect of the i^th^ marker, and ***u*** is a vector of genetic values, modelled as 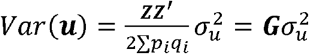 where ***G*** is a genomic relationship matrix and **Z** is a matrix with gene contents for all markers; *p*_*i*_ and *q*_*i*_ are the frequencies of the major and minor alleles of the i^th^ marker, respectively. We assume that variances 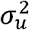 and 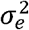 are known with high certainty, which is usually the case in large data sets. The normal test for the effect of the marker uses the statistic 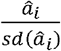, where both values are obtained from inversion of the linear equations of the model. P-values are obtained as 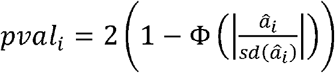 where Φ is the cumulative standard normal function.

It has been shown [23–25] that joint estimates of marker effects ***α*** from SNP-BLUP (or GBLUP models) of the form ***y*** = ***Xb*** + ***Za*** + ***e*** with prior assumption 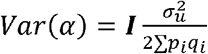 lead themselves to statistics 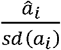 that are equivalent to the statistic obtained for the same marker in the “fixed regression” framework described above. The application of SSSNP-BLUP instead of SNP-BLUP is straightforward, because 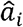 and 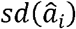 are immediately available (e.g. [15]). In GBLUP and SSGBLUP, the same values can be obtained as linear transformations of the solutions for genetic values 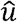 and their Prediction Error (Co)Variances [23–26]. These can be obtained from the inverse of the Mixed Model Equations[28,29].

### Algorithm

The algorithm for SSGWAS, which includes genotyped and non-genotyped animals, has been implemented in the blupf90 suite of programs [30] as follows. It combines the algorithms for SSGBLUP (e.g.[16]) and back-solving to obtain marker estimates and their associated p-values from estimates of breeding values [25]:

1. Construct the joint pedigree-genomic relationship matrix 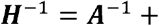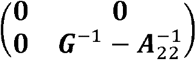, that is the inverse of 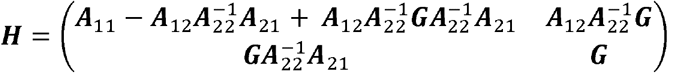, which projects genomic relationships ***G*** = ***ZZ***′/2∑*p*_*i*_*q*_*i*_ from genotyped animals (labelled as “2”) to non-genotyped animals (labelled as “1”). Matrix 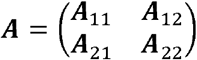 is the pedigree-based relationship matrix. Matrix **G** is (usually) constructed as 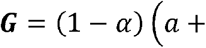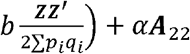 with *a* and *b* obtained to equate average inbreeding and average relationships in ***G*** and ***A***_22_, and *α* a small value (typically from 0 to 0.05). In this way, both relationships are “compatible” [31,32] and ***G*** is invertible [10]. Matrix **Z** contains centered gene content as in [10] but using observed allele frequencies.
2. Construct the mixed model equations for SSGBLUP. In a simple case (a model with a single genetic effect) this would be:

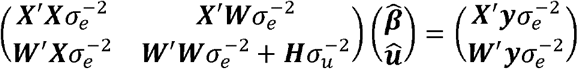

Where ***β*** are “fixed” effects and **u** are individual animal (not marker) effects.
3. Factorize and obtain the sparse inverse of the coefficient matrix. The whole inverse cannot be obtained directly as it is typically too big [33]. Most exactly, a “sparse inverse” (***C***) is obtained with selected elements of the inverse corresponding to the non-zero entries of a (sparse) Cholesky factor (***LL***′). In our case, this is achieved using supernodal sparse factorization and inversions as programmed in YAMS [34,35]. Factorization is the computing bottleneck of the procedure and is roughly cubic on the number of genotyped animals; YAMS reduces the computing time by, roughly, one order of magnitude.
4. Solve the mixed model equations for 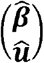 using the sparse Cholesky factor.
5. Extract from ***C*** the submatrix corresponding to the genotyped animals, 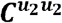. This matrix contains the prediction error (co)variances of their estimated breeding values, 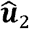, i.e. 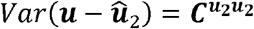.
6. Backsolve for SNP effect estimates:

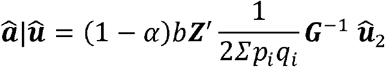

If matrix **G** is full rank and compatible with pedigree relationships (for instance, if ***Z*** is built with base allele frequencies) then 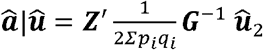 [10,36,37].
7. Obtain individual prediction error variances of SNP effects as [25]

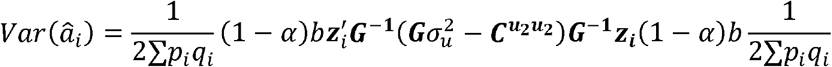

where ***z***_*i*_ is the i-th column of matrix **Z**, corresponding to genotypes of marker *i* across individuals. The values of 0.95 in steps 7 and 8 refer to the “blending” of **G** with **A** in step 1 and it will change with different choices of blending parameters.
8. Finally, the p-value for marker *i* is obtained as 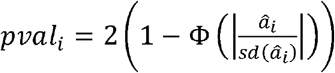.

Note that this analysis has to be run only once, as opposed to the *n* individual runs for *n* markers in the classical “fixed effect” regression or EMMAX (e.g. [38]).

## MATERIAL

We re-analyzed a data set from the American Angus Association [39] including birth weight. As the complete data set is very large, with about 7 million individual weights, 52000 genotyped animals and 8 million animals in pedigree, we used only partial data. These included phenotypes recorded in the last four years, specifically from 2009 to 2012, which comprised 1,046,623 birth weights. Three generations of ancestors were traced back, totaling 1,849,865 individuals. All available genotypes of sires with phenotyped offspring were used (i.e., 1,424 genotyped sires). Genotypes were obtained with the BovineSNP50k v2 BeadChip; 38,122 polymorphic markers were used after quality control [39].

The linear model for birth weight included the effects of contemporary group, animal additive genetic effect, and a permanent environmental effect of the mother, which considers the maternal ability during pregnancy. This differs from the model actually used in national evaluations, that also considers a maternal genetic effect. Variance components were fixed at values from the national evaluation, with heritability of 0.48 and a maternal component of 0.10.

Computations were done using the blupf90 software suite. Then, GWAS results were plotted with qqman [40]. Rejection thresholds used a Bonferroni correction for multiple testing of 0.05/38122, which equals to 5.9 in the −*log*10 scale. Significant regions were explored in AnimalQTLdb and Jbrowse [41] using the bovine map assembly UMD 3.1 [42]. Although a new genome map assembly has already been published (i.e., ARS-UCD 1.2), the aforementioned genome browsers are still using the UMD 3.1.

In addition to exact SSGWAS using p-values, we also ran “approximated” SSGWAS using estimates of marker effects, in the form of percentage of variance explained by marker effect [18]. This estimates the population genetic variance explained by the marker effect, and is approximately computed as 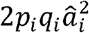. There are no theoretical thresholds in this approach, and we used an arbitrary threshold of 0.10% of total genetic variance explained by 1 marker. Opposite to [18], we do not present results of iterating the SSGBLUP using “weights” for each marker, as this procedure did not result in increased marker effect and variance (results not shown).

## RESULTS

Factorization of the Mixed Model Equations and extraction of 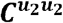 required 30Gb of RAM memory and the computing time was 14h (wall-clock time). Computation of 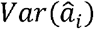 and p-values took a negligible time of a few minutes. Quantile-quantile plot (QQplot) and Manhattan plots are in Figures 1 and 2. The QQplot does not show large deviations from the null hypothesis, which means that SSGBLUP captures correctly the structure of the population *via* relationship matrices. In presence of structure unaccounted for, inflation of p-values is expected [3,43,44].

The GWAS results pointed two regions significant at genome-wide level, with values of −*log*10(*pvalue*) near 8. The first region at the end of chromosome 7 includes 3 markers: ARS-BFGL-NGS-107035, ARS-BFGL-NGS-101886 and ARS-BFGL-NGS-18900, which are all in very high linkage disequilibrium (correlation > 0.95 in all cases). Very close signals in the same region can be found in *animalQTLdb* for the trait “body conformation trait” in Holstein cattle [8] and “average daily body weight gain" found in Brangus (composite of Brahman and Angus) [45]. The fact that the signal appears in several breeds would imply that this is not a false positive. The second region at the end of chromosome 20 only includes marker Hapmap42635-BTA-68718. This marker is included in a QTL region detected by [46] in Hereford.

**FIGURE 1.**
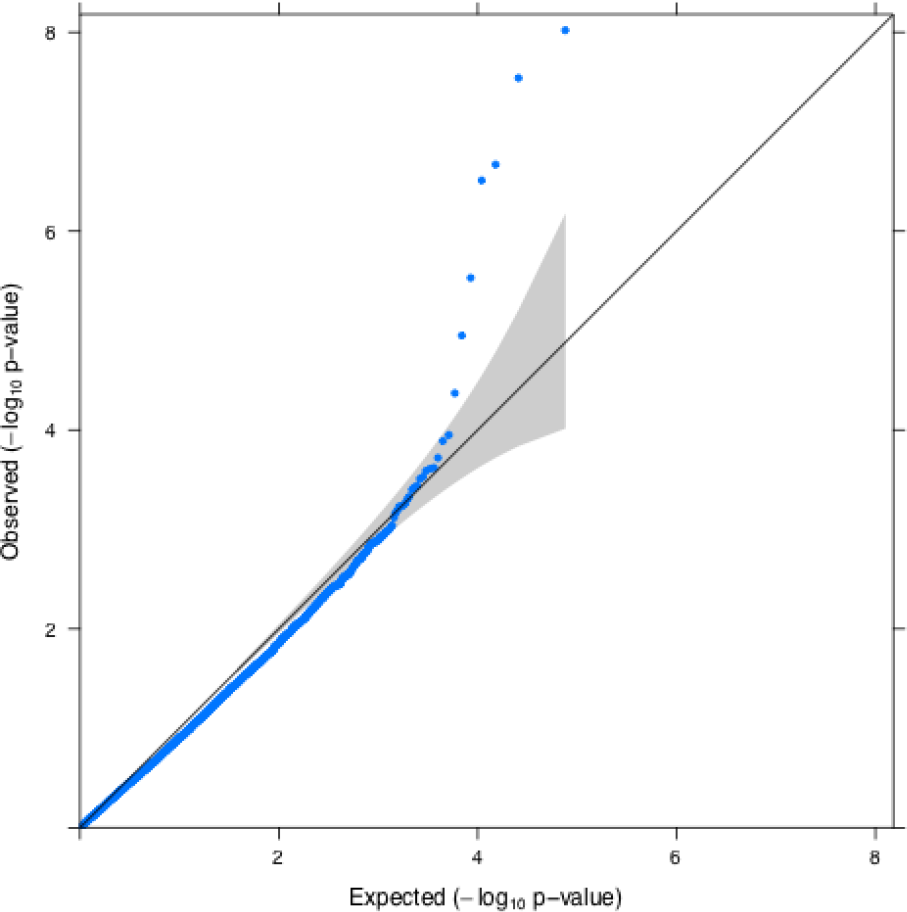
Quantile-quantile plot (QQPLOT) for the The grey region represents a 95% confidence interval.

**FIGURE 2.**
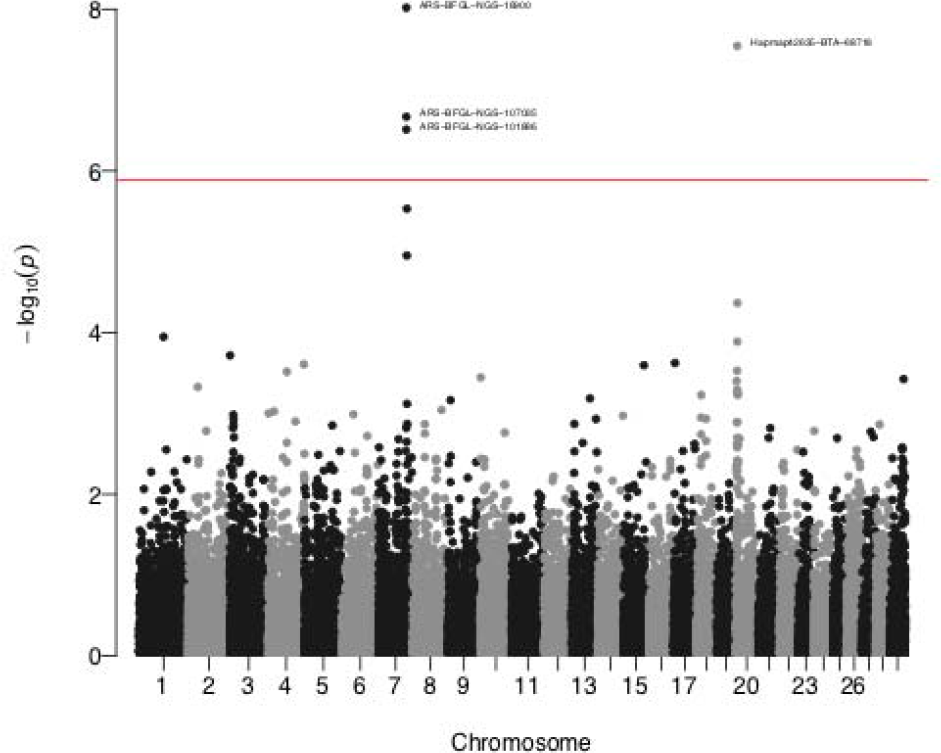
Manhattan plot (p-values of individual SNP effects) for birth weight. The red line corresponds to Bonferroni rejection threshold for nominal alpha=0.05.

Figure 3 shows the percentage of genetic variance attributed to each marker. The top markers are the same as in Figure 2 (ARS-BFGL-NGS-18900 and Hapmap42635-BTA-68718), but the region in chromosome 7 has a shorter peak than the region in chromosome 20. This is at first sight perplexing, because estimates of marker effects and standard errors are similar for peaks at chromosomes 7 and 20 (effects ± standard deviations of −0.041±0.007 and 0.043 ± 0.008, respectively). The different height of the peaks in chromosome 7 and chromosome 20 in the variance plot (Figure 3) and in the −*log*10(*pvalue*) plot (Figure 2) is entirely due to the different minor allelic frequencies: 0.28 and 0.46 respectively.

**FIGURE 3.**
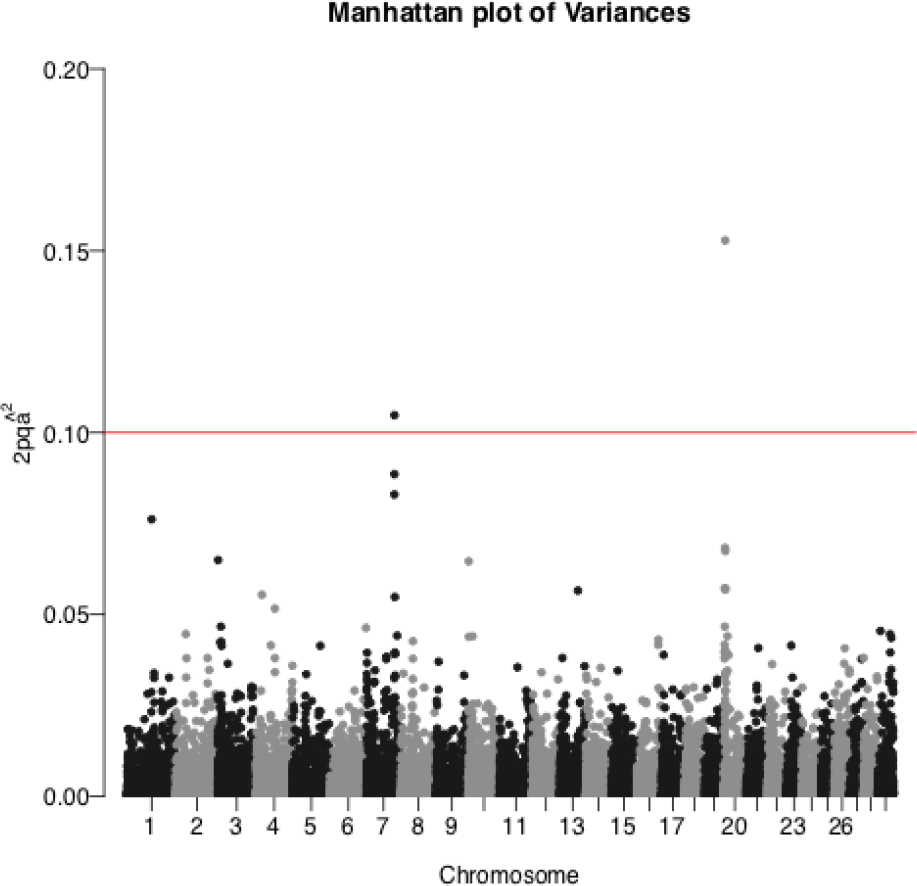
Percentage of genetic variance explained by markers for birth weight in American Angus. The red line corresponds to an arbitrary rejection threshold of 0.10%.

## DISCUSSION

Estimates of SNP effects from single-step methods became available in 2012 [18,47]; however, a measure of significance for SNP estimates has not been available until now in current implementations of single-step GBLUP. Expanding the ideas from the equivalence between GBLUP and GWAS [23–25], Lu et al. [27] derived exact GWAS for a single-step GBLUP framework in a relatively small data set. In our study we showed exact p-values calculated for the first time using a large data set including genotypes, phenotypes, and pedigree.

Known associations from beef cattle were used to validate the proposed methodology and the results show good agreements with the literature. The qqplot showed no inflation of p-values, as expected, because the relationship structure is well accounted for [2,3,43]. Additional advantages compared to “approximate” SSGWAS include avoiding the need for arbitrary choices as the length of segments, percentage of variance explained, and iterative scheme. The use of percentage of variance explained, as advocated by several authors [18,22], is misleading because it depends too much on the heterozygosity at the marker. Variance explained may be useful for breeding purposes, but may not be useful for QTL detection, where the final objective is a detailed understanding of gene action.

Although SSGWAS was possible using the last 4 years of phenotypes and all genotyped bulls present in the American Angus data, the computational burden for the factorization of the left-hand side of the mixed model equations was not negligible. In our case, matrix **H**^−1^ contains (roughly) 16 million non-null elements from pedigree and 2 million non-null elements from genotypes. However, the matrix **H**^−1^ for the whole data set in Lourenco et al (2015) contained 63 million non-null elements from pedigree and 2704 million non-null elements from genotypes. The first simple method to reduce the size of the problem is to include only genotyped ancestors in exact SSGWAS and exclude candidates to selection. This leads to a number of genotyped animals in the thousands or tens of thousands. The second strategy is to use the sparse version of **G**^−1^ known as APY [39,48,49] that enormously increases the number of null elements in **G**. If animals in the core portion of **G** are a representative sample of the population, this also improves the estimates of the SNP effects [50].

We emphasize that, contrary to regular GWAS, in SSGWAS computations do not need to be repeated at each marker, therefore, all p-values were obtained in a single run of SSGWAS.

## CONCLUSIONS

The exact Single Step GWAS is a very general and efficient strategy for QTL detection, localization and testing, providing accurate p-values. It can be used in complex data sets such as the ones in animal breeding, with a host of unbalanced effects, very complex models and the presence of genotyped and ungenotyped animals. Our proposed strategy is computationally viable for very large populations and solves the main issues in the current Single Step GWAS, that were precluding the use of the method.

## DECLARATIONS

### ETHICS APPROVAL AND CONSENT TO PARTICIPATE

Not applicable.

### CONSENT FOR PUBLICATION

Not applicable.

### AVAILABILITY OF DATA AND MATERIALS

The data that support the findings of this study were provided from the American Angus Association but restrictions apply to the availability of these data, which were used under license for the current study, and so are not publicly available. The methods described here are included using “OPTION snp_p_value” in the parameter file in software blupf90 (factorization of the mixed model equations and solving of the SSGBLUP equations) and postGSf90 (backsolving of snp effects and computation of p-values), available at http://nce.ads.uga.edu/software/.

## COMPETING INTERESTS

The authors declare that they have no competing interests.

## FUNDING

This study was partially funded by the American Angus Association (St. Joseph, MO) and by Agriculture and Food Research Initiative Competitive Grants no. 2015-67015-22936 from the US Department of Agriculture’s National Institute of Food and Agriculture.

## AUTHORS CONTRIBUTIONS

The theory was refined by IA, AL and FC. IA and YM programmed the algorithms. The American Angus data set was analysed by AL and DL. DL and IM contributed critically to intperpretation of results and planning of the anlyses. The manuscript was initially written by AL and then completed by the rest of authors. All authors read and approved the final manuscript.

## ACKNOWLEDGMENTS

We acknowledge Dan Moser and Stephen Miller for discussions, corrections to the manuscript, and for sharing the American Angus Association data through the Angus Genetics Inc. Discussions with Anne Ricard (IFCE, Toulouse, France) and Toni Reverter (CSIRO, Brisbane, Australia) are highly appreciated. AL did part of this work while visiting the University of Georgia in a four-month sabbatical.

